# Transcription-factor centric genome mining strategy for discovery of diverse unprecedented RiPP gene clusters

**DOI:** 10.1101/467670

**Authors:** Shaozhou Zhu, Guojun Zheng

## Abstract

Ribosomally synthesized and post-translationally modified peptides (RiPPs) are a rapidly emerging group of natural products with diverse biological activity. Most of their biosynthetic mechanisms are well studied and the “genome mining” strategy based on homology has led to the unearthing of many new ribosomal natural products, including lantipeptides, lasso peptides, cyanobactins. These precursor-centric or biosynthetic protein-centric genome mining strategies have encouraged the discovery of RiPPs natural products. However, a limitation of these strategies is that the newly identified natural products are similar to the known products and novel families of RiPP pathways were overlooked by these strategies. In this work, we applied a transcription-factor centric genome mining strategy and diverse unique crosslinked RiPP gene clusters were predicted in several sequenced microorganisms. Our research could significantly expand the category of biosynthetic pathways of RiPP natural products and predict new resources for novel RiPPs.

## 1 Introduction

Intensive abuse of antibiotics for human therapy, farm animals, and fish in aquaculture has resulted in the occurrence of multiresistant pathogens[1]. Currently, it is a growing public concern, as it is associated with increased number of deaths and suffering for humans. The multidrug resistant strain, *Staphylococcus aureus*, is one of the most common pathogens that cause serious diseases[2, 3]. Accelerating the development of novel antibiotics with novel antimicrobial mechanisms is still the major strategy to treat these potentially life-threatening infections[4, 5].

Microbes are still the richest source of natural products and are used for the discovery of new antibiotics[6]. For example, newly developed antibiotics such as teixobactin and lassomycin have been derived from soil bacteria[7]. Generally, most of these antibiotics are assembled by multimodular megasynthases, like nonribosomal peptide synthetases (NRPS) and polyketide synthases (PKS) or NRPS-PKS hybrids[8-11]. In addition, ribosomally synthesized and post-translationally modified peptides (RiPPs) are becoming a major group of antibiotics, which were underestimated in the past[12]. With the rapid progress of genome sequencing technology, many of the RiPP biosynthetic gene clusters have been uncovered through diverse genome mining strategies[13-15]. Unlike the broad-spectrum antibiotics, most RiPPs with antibiotic activity often show a very narrow activity spectrum, which can benefit patients, as their side effects will be lesser than the broad-spectrum antibiotics[14, 16].

In general, the biosynthesis of RiPPs occurs in a ribosome-dependent manner[12]. A precursor peptide with a length of 20-100 proteinogenic amino acids was first assembled on the ribosome. Typically, the precursor peptide is divided into two parts: An N-terminal leader sequence and a C-terminal core sequence[12, 15, 16]. The leader peptide is used to guide the biosynthesis and is subsequently removed by a specific transporter or a peptidase. With the assistance of the leader peptide, extensive post-translational modifications of the core peptide are performed by the tailoring enzymes, often within close proximity to the precursor gene[17]. With this strategy, although the precursor peptide relies on the limited proteinogenic amino acids, structurally diverse compounds were created[12]. Common structural features of RiPPs include methylations, phosphorylation, heterocycles, backbone crosslinks, dehydrated amino acids, and many others.

To find novel antibiotics, a couple of strategies have been developed[18-21]. Among them, bioinformatics strategy is the most powerful method because of its culture independent fashion[13, 15, 18]. Moreover, in some cases the final structures of the compounds produced by the identified biosynthetic gene clusters could be predicted, which helps reduce the rediscovery of known compounds. Currently, most bioinformatics strategies developed depend on domain homology[13, 15, 18]. Of all the existing biosynthetic gene cluster-predicting tools, the most popular one is antiSMASH, which is widely used for the prediction biosynthetic pathways of secondary metabolites[22, 23]. Generally, the genes for the biosynthesis of natural products are organized in clusters. This unique feature laid the foundation for the development of these bioinformatics tools. By collecting the protein domains found in known clusters these tools could use the database to predict new clusters by searching for similar domains[22, 23]. This genome mining strategy is very effective for the discovery of similar clusters, but cannot predict completely new biosynthetic gene clusters with novel proteins containing unknown domains.

As for the RiPP family of natural products, most of the known RiPP biosynthetic gene clusters are well studied[12, 16]. The co-existence of the precursor peptide and the specific tailoring enzymes ultimately define the classes of RiPPs. Past research has shown that bioinformatics studies could readily identify novel RiPPs using homology searching methods[15]. For example, novel modified lasso peptides, such as phosphorylated and acetylated lasso peptides, were discovered using this strategy[24-26]. However, completely new families of RiPPs have not been discovered by these strategies, which hinders the progress of RiPP research. To develop novel bioinformatics strategies for the identification of completely new families of natural products, we focus on transcription factors that are important markers for the biosynthesis of natural products. Our strategy is based on the idea that different biosynthetic pathways could be co-regulated by a similar regulator.

Here, we present a transcription-factor-centric genome mining strategy. We also searched for potential novel RiPP gene clusters in genome-sequenced microorganisms. Through bioinformatics analysis and precursor prediction, we have identified diverse families of RiPP biosynthetic pathways. The results showed that several microbes possess the potential to synthesize a variety of unprecedented, crosslinked, ribosomally synthesized, and post-translationally modified peptides.

## 2 Results

### 2.1 Selection of the SHP/Rgg quorum sensing system as an inquiry for genome mining

To find a proper transcription-factor for genome mining studies, we search for a potential proper quorum sensing system and the streptococcal bacteria draws our attention. Research has shown that these bacteria use a novel quorum sensing system for intraspecies communication and regulate the biosynthesis of natural peptide products[27, 28]. One example is the recently identified peptide-based quorum sensing system from *Streptococcus thermophilus*, which is a non-pathogenic bacterium widely used in the fermentation of dairy products[29-31]. They have shown that *S. thermophilus* can produce a short hydrophobic peptide (SHP, H2N-EGIIVIVVG-COOH) that functions as a pheromone[29-31]. Generally, the peptide is exported into the environment after synthesis. However, when the concentration of SHP in the environment reaches the threshold, it triggers the Ami oligopeptide transporter and is imported into the cytoplasm again. SHP then binds to the transcription factor Rgg and regulates diverse cellular processes, including the biosynthesis of streptide (Fig. 1)[29-31]. Streptide is a new family of RiPPs that are biosynthesized via a short pathway containing three genes: *strA*, which encodes the precursor peptide, *strB*, a radical S-adenosylmethionine (SAM) enzyme, and *strC*, an ABC transporter[32-35]. The strB enzyme is responsible for the formation of the unusual lysine-to-tryptophan crosslink. *strC* is needed to remove the leader peptide and export the mature streptide outside the cell. The discovery of streptide, and the elucidation of its biosynthesis pathway is an impressive example of the biochemical pathways catalyzed by the radical SAM enzymes[32-35]. Past research on sactipeptide biosynthesis also showed that the thioether bond formation was catalyzed by similar radical SAM enzymes[36]. All these examples demonstrate that they are one of the most versatile enzyme families and are widespread in nature. Therefore, searching for novel radical SAM enzymes containing RiPP gene clusters could present opportunities for finding novel natural products with unique biosynthetic reaction mechanisms.

**Fig. 1.**
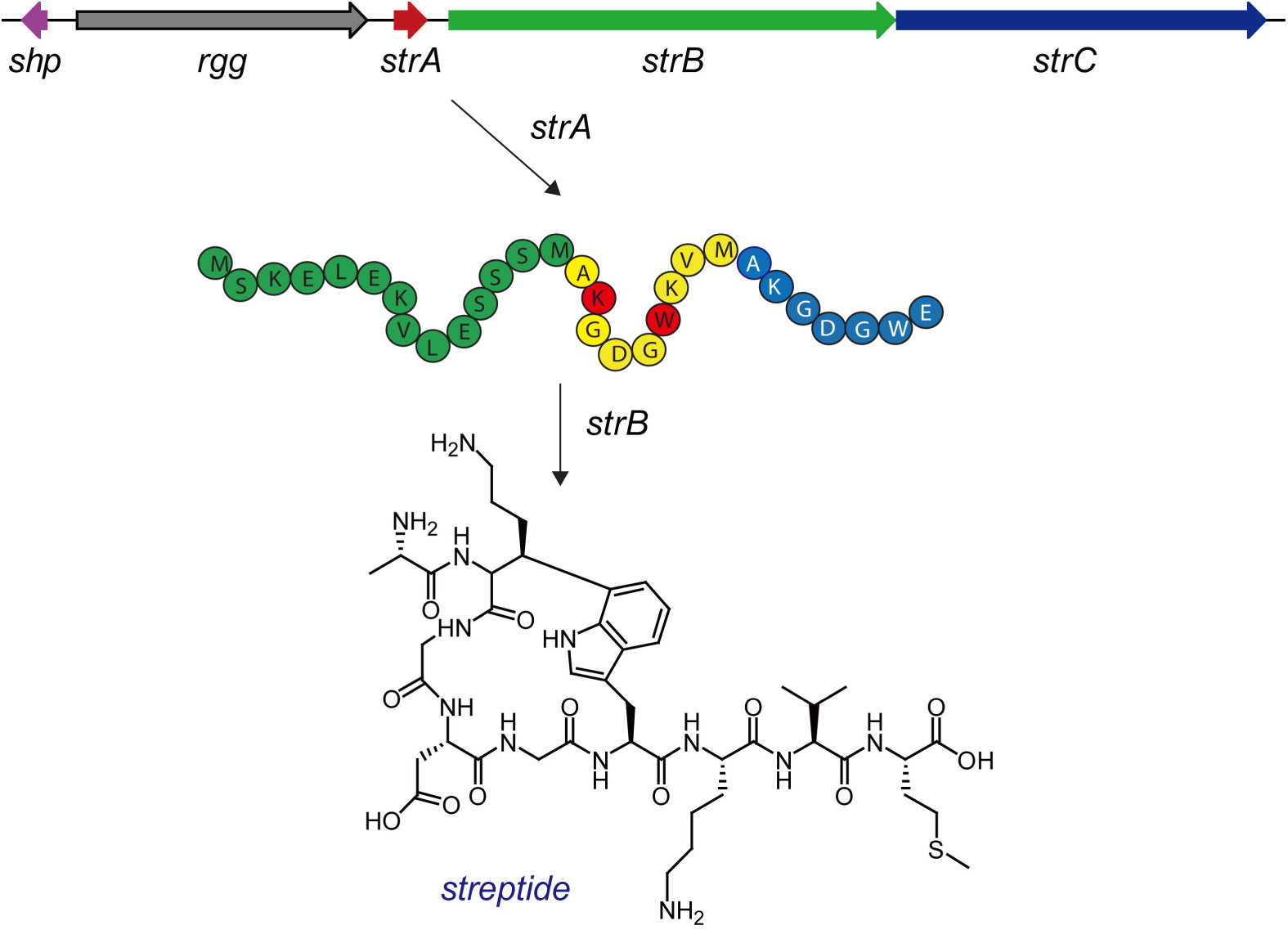
Representative structure and biosynthesis of streptide: A unusual lysine-to-tryptophan crosslink in strA was installed by a Radical SAM enzyme strB. The biosynthetic pathway was regulated by the shp/Rgg quorum sensing system.

The SHP/Rgg quorum sensing system has been shown to regulate diverse cellular processes[29]. We were curious about the distribution of these systems in other bacteria. One hypothesis was that such kind of transcription factors might control different types of RiPP pathways. This led us to develop a transcription-factor centric genome-mining approach for the discovery of novel biosynthetic pathways. To confirm our hypothesis, the Rgg gene (WP_011681385.1) from *S. thermophilus* was selected as a query sequence and a PSI-BLAST (position-specific iterated BLAST) was performed using the NCBI (national center for biotechnology information) database[37]. The algorithm parameters were set as follows: Maximum target sequences to display was set to 500; expect value threshold was set to 10; BLOSUM62 was chosen as the matrix for scoring parameters. Surprisingly, this bioinformatics survey indeed showed that a similar type of quorum sensing system widely exists in nature, especially in the streptococcal bacteria. Further analysis showed that the loci were upregulated by these quorum-sensing systems, which lead to the discovery of a large number of novel RiPP biosynthetic pathways. Here, we will show three different representative families of novel RiPP biosynthetic pathways identified by this strategy and the structure prediction of the final natural products.

### 2.2 The WGK type of RiPP biosynthetic pathways

By the genome mining study on Rgg from *S. thermophilus*, 500 new Rgg regulators were identified. After the identification of these novel transcription-factors, the associated biosynthetic pathways present downstream were studied in detail. Surprisingly, this strategy turned to be very successful and several new RiPP biosynthetic pathways were discovered. The first apparent new family of RiPP gene clusters included those from *Streptococcus mutans* ATCC25175, *Streptococcus equi subsp. Zooepidemicus H70*, *Streptococcus equinus* strain Sb09, *Streptococcus ferus* DSM 20646, *Streptococcus sp*. 45, and *Enterococcus caccae* ATCCBAA-1240. In those systems, three genes were directly located downstream of the Rgg like transcription factors (Table 1). Like the streptide system, the genes are a radical SAM enzyme and an ABC transporter. Besides, there is a PqqD domain protein adjacent to the radical SAM enzyme (Fig. 2A). This domain is observed in several RiPP biosynthetic pathways. In the biosynthesis of pyrroloquinoline quinone (PQQ), the PqqD was shown to assist the tailoring enzyme PqqE and crosslink a tyrosine residue to a glutamate residue in an intramolecular cyclization of PqqA[38, 39]. In the biosynthesis of lasso peptides, such as the paeniondin and streptomonomicin, the PqqD domain was shown to bind to the leader peptide and transfer its peptide substrate to other enzymes for processing[40, 41]. Apart from these examples, the PqqD domain (precursor peptide recognition element) is also present in other diverse RiPP biosynthetic pathways, such as linear azole-containing peptides, thiopeptides, and sactipeptide[40]. It is an important marker for novel RiPP biosynthetic pathways, since it is used in leader peptide binding. Thus, these gene clusters belong to a novel RiPP biosynthetic pathway.

**Fig. 2.**
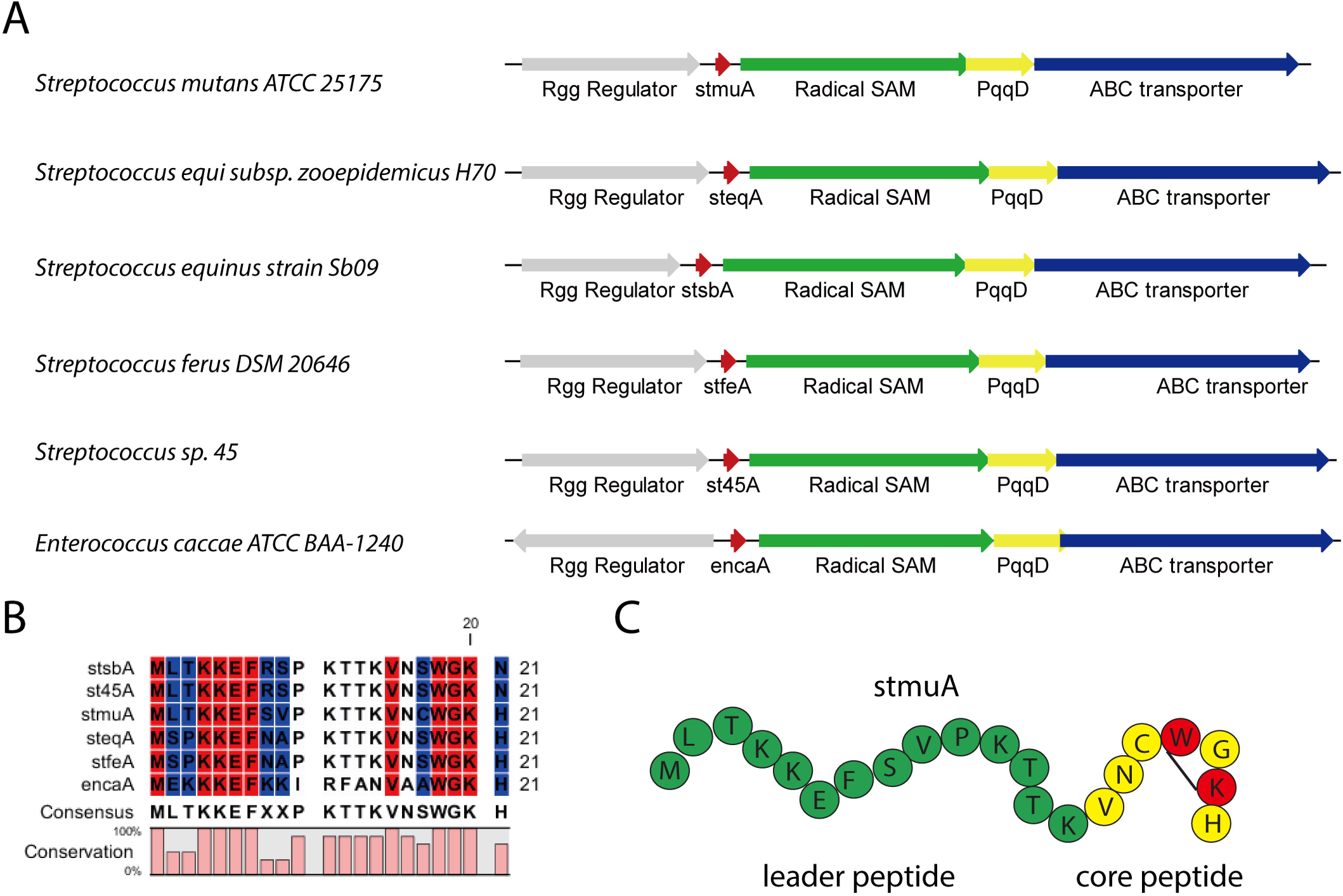
The WGK type of RiPP biosynthetic pathways (A) Organizations of six identified such kind of biosynthetic gene clusters. Gene products marked in colar: Red (precursor peptide), Green and yellow (Radical SAM enzymes and PqqD domain), blue (ABC transporter) (B) Alignment of the potential precursor peptides. (C) Precursor peptide colors of stmuA: leader peptide (green), core peptide (yellow), potential crosslink was marked in black.

**TABLE 1.**
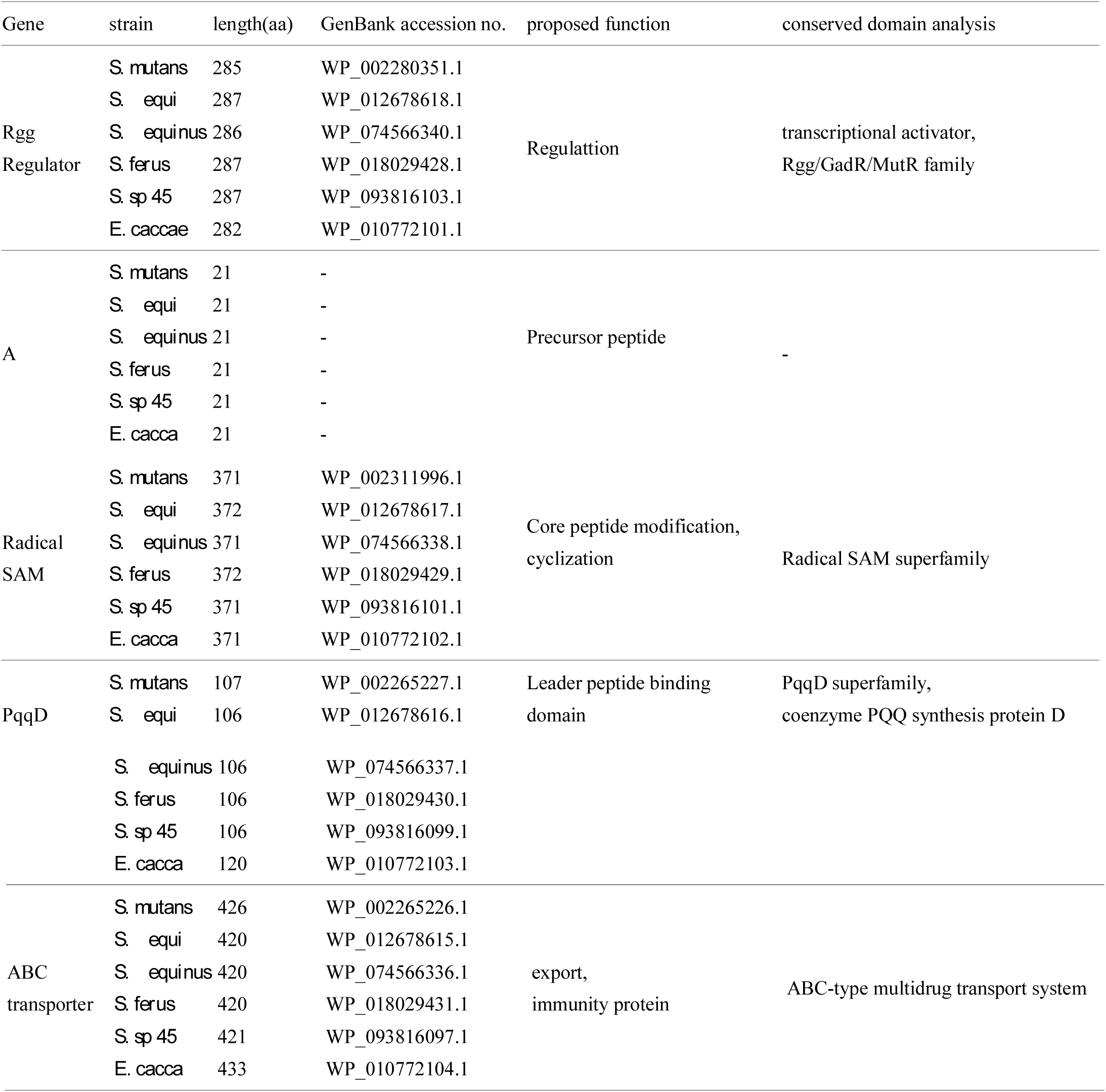
Proposed functions of the ORFs in the WGK type of RiPPs biosynthetic pathways.

The sequences of all these identified gene clusters were downloaded from NCBI and analyzed by Clone Manager for target precursor peptides. Usually, the precursor peptide is located in front of the processing enzymes. In the case of streptide, the precursor peptide was located between the Rgg Regulator and the strB protein[33]. The precursors of these newly identified pathways likely follow the similar synthetic logic. Therefore, we looked for short ORFs in this area and a single potential precursor peptide was identified in each cluster in this region. The sequences of these precursor peptides are shown in Fig. 2B and were analyzed using the CLC Main Workbench. All the precursor peptides consist of 21 aa. Based on the alignment, these peptides have two obvious conservation regions, which likely consist of the leader peptide and the core peptide. The conserved reside, KKEF, in the leader peptide region is probably used to bind to the PqqD domain. On the other hand, the WGK conserved in the core peptide region might contribute to the final structure of the RiPP natural products. We hypothesized that there might be an unusual cyclization between these residues catalyzed by the radical SAM enzymes and the PqqD domain (Fig. 2C). Further *in vivo* or *in vitro* studies on these systems are needed to confirm our hypothesis. Nevertheless, considering the sequence of the precursor peptide and the components of the biosynthetic enzymes, these newly identified RiPP biosynthetic pathways are distinct from the streptide systems. Thus, they represent a completely new family of RiPP gene clusters.

### 2.3 The CGPSHSCGGGR type of RiPP biosynthetic pathways

After we discovered the WGK-type RiPP biosynthetic pathway, we further studied the identified transcription factors and another new types of RiPPs were identified and classified. These gene clusters are from *Streptococcus thermophilus* JIM 8232, *Streptococcus constellatus* subsp. pharyngis C232, *Streptococcus gallolyticus* DSM 16831, *Streptococcus gordonii* strain G9B, *Streptococcus himalayensis*, and *Streptococcus parasanguinis* CC87K (Fig. 3A). In those systems, three genes were located downstream of the Rgg like transcription factors including a short precursor peptide (Table 2). Unlike the previously identified WGK type of RiPP biosynthetic pathway, where a PqqD domain was present, these gene clusters are more similar to the streptide system[33, 35]. A radical SAM enzyme and an ABC transporter could be found downstream of the Rgg Regulator. In five cases, except the one from *S. himalayensis*, the precursor peptides were predicted using the NCBI database. They were directly located between the regulator and the process enzyme.

**Fig. 3.**
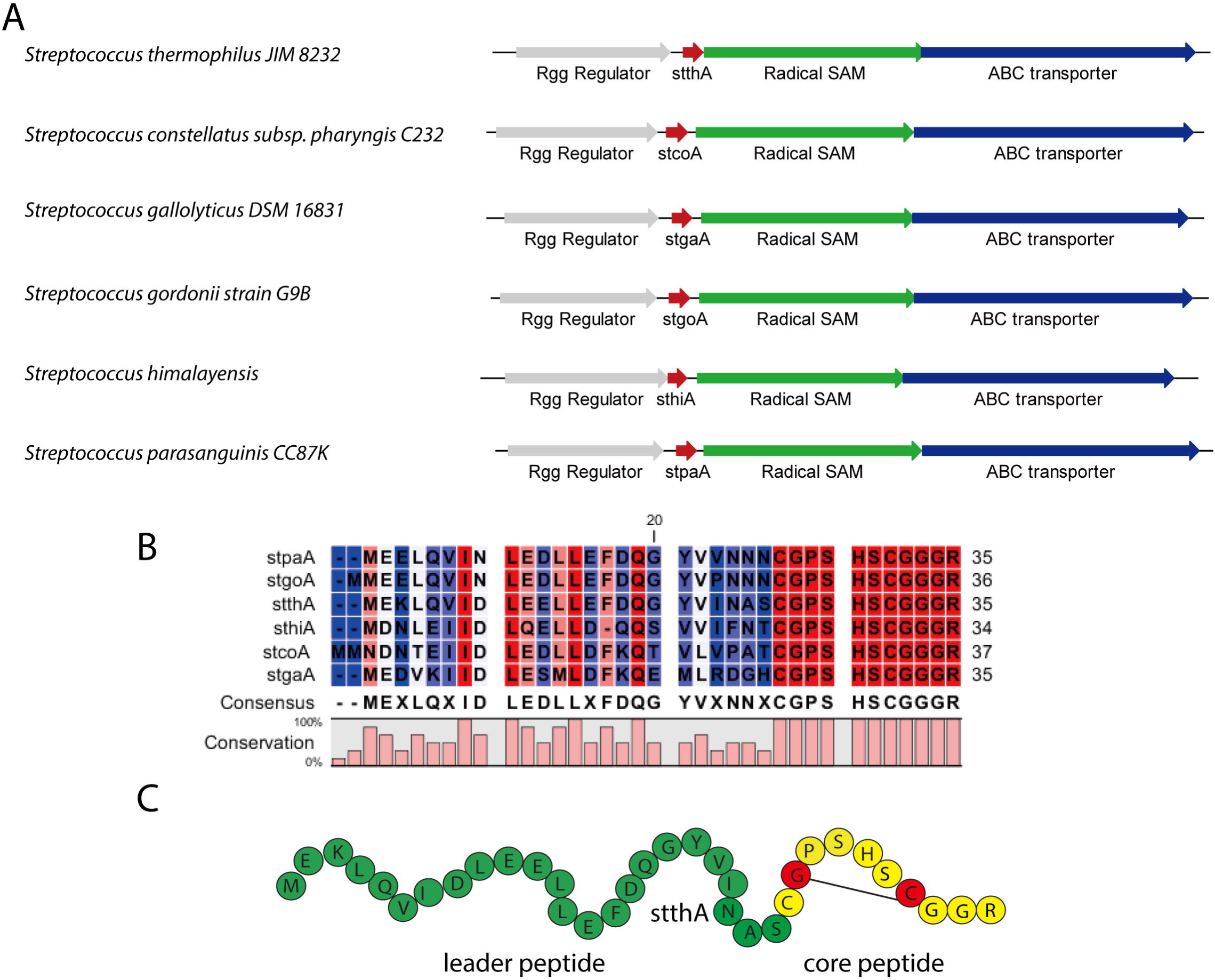
The CGPSHSCGGGR type of RiPP biosynthetic pathways (A) Organizations of six identified such kind of biosynthetic gene clusters. Gene products marked in colar: Red (precursor peptide), Green (Radical SAM enzymes), blue (ABC transporter) (B) Alignment of the potential precursor peptides. (C) Precursor peptide colors of stthA: leader peptide (green), core peptide (yellow), potential crosslink was marked in black.

**TABLE 2.**
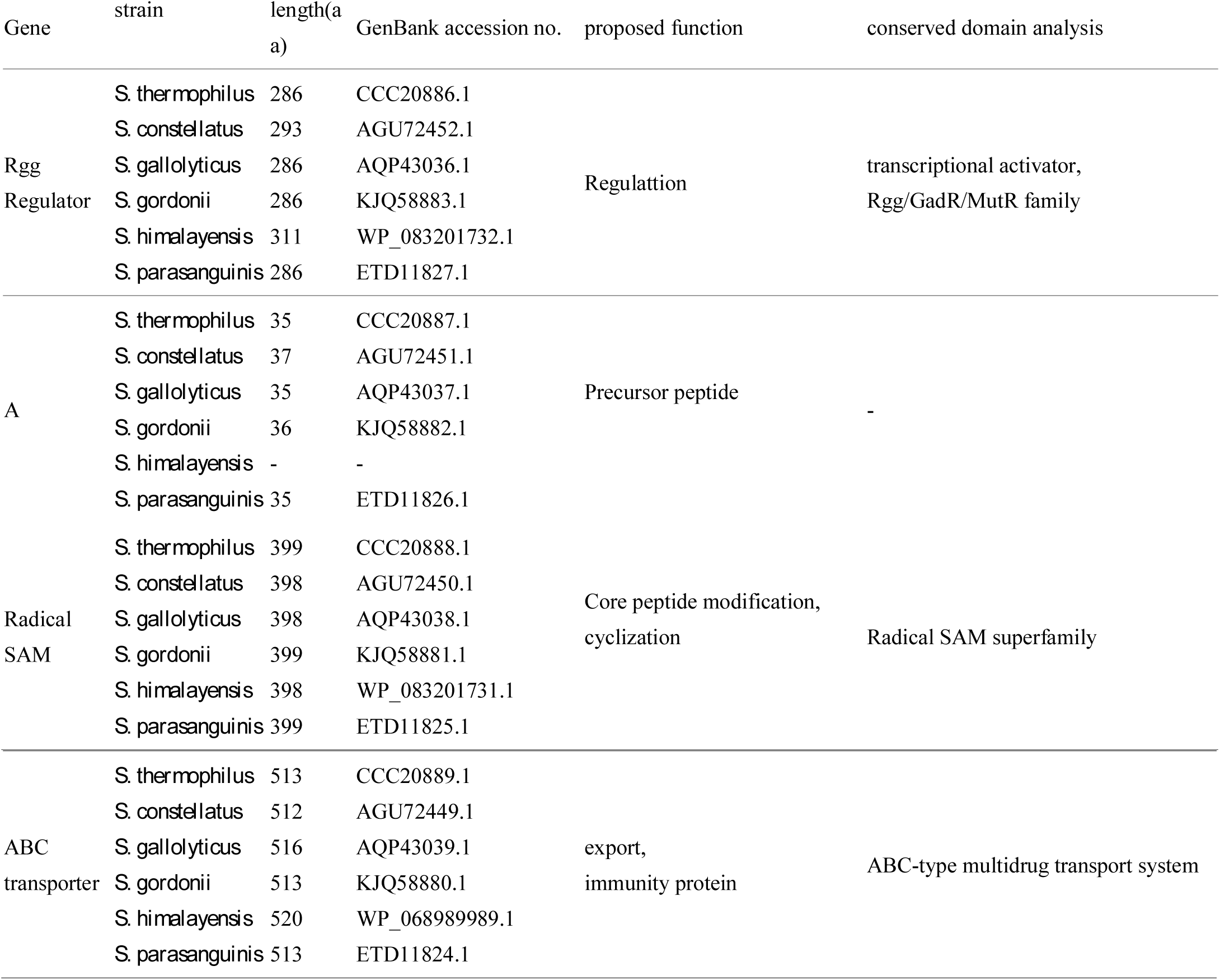
Proposed functions of the ORFs in the CGPSHSCGGR type of RiPPs biosynthetic pathways.

The sequence of all these identified gene clusters was downloaded from NCBI and the precursor peptide from *S. himalayensis* was also predicted based on short-ORF searches. These six precursor peptides are shown in Fig. 3B. They were analyzed using the CLC Main Workbench. These precursors have a length of about 34-37 aa. Based on the sequence alignment, these potential precursor peptides have a highly conserved region at the center of the N terminus, which might be the binding site for the process enzymes. The C terminus is also highly conserved and the sequences are the same in all six cases. This part consists of the core peptide and has a conserved sequence “CGPSHSCGGGR”. We also hypothesized that the radical SAM enzyme will introduce an unusual modification at that region and the leader peptide will be removed by the ABC transporter (Fig. 3C). Although, the biosynthetic mechanism of these systems is similar to that of the streptide system, the distinct precursor peptide and a dissimilar radical SAM enzyme make these systems another completely new family of RiPPs.

### 2.4 The ITRRRY type of RiPP biosynthetic pathways

The third type of system discovered based on this transcription-factor centric genome-mining approach is observed in *Staphylococcus pseudintermedius*, *Streptococcus parauberis*, *Streptococcus* sp. UMB0029, and *Streptococcus suis*. Downstream to the Rgg type regulator, two enzymes could be found. This system also has a radical SAM enzyme and an ABC transporter similar to the streptide system and the previous CGPSHSCGGGR type of RiPP system[33, 35]. However, the radical SAM enzyme showed a very low similarity compared to the one from the streptide biosynthetic pathway, indicating that they are also a completely new family of RiPP biosynthetic pathway. All these gene clusters were downloaded from NCBI and the accession numbers are shown in Table 3. The potential precursor peptides were detected between the Rgg regulator and the radical SAM enzyme by searching for short ORFs in this region (Fig. 4A). Four potential precursor peptides were predicted and an alignment was performed using the CLC Main Workbench. As shown in Figure 4B, these precursor peptides also are clearly divided into two parts. The N terminus has a conserved sequence, which contains the leader peptide and is likely involved in the binding process of the enzymes. On the other hand, the C terminus has a conserved sequence, namely “ITRRRY”, which contributes to the final products. These systems are named as the ITRRRY type of RiPP. Like the streptide system, we hypothesized that the radical SAM enzymes might catalyze an unusual cyclization reaction in the core peptide (Fig. 4C). Since these are new classes of natural products, the prediction of the final structure is difficult and further *in vivo* and *in vitro* studies are necessary to confirm our hypothesis.

**Fig. 4.**
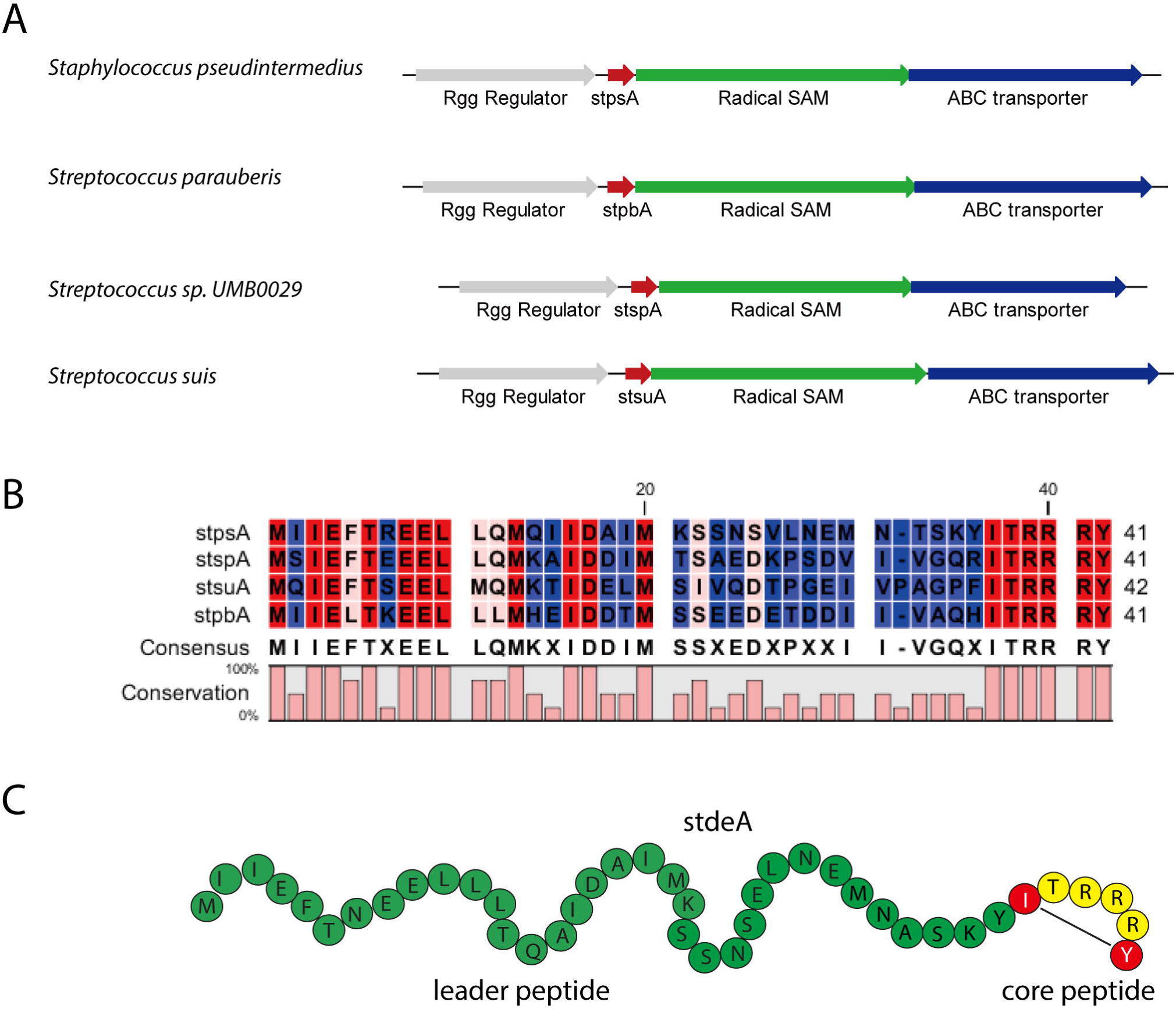
The ITRRRY type of RiPP biosynthetic pathways (A) Organizations of four identified such kind of biosynthetic gene clusters. Gene products marked in colar: Red (precursor peptide), Green (Radical SAM enzymes), blue (ABC transporter) (B) Alignment of the potential precursor peptides. (C) Precursor peptide colors of stdeA: leader peptide (green), core peptide (yellow), potential crosslink was marked in black.

**TABLE 3.**
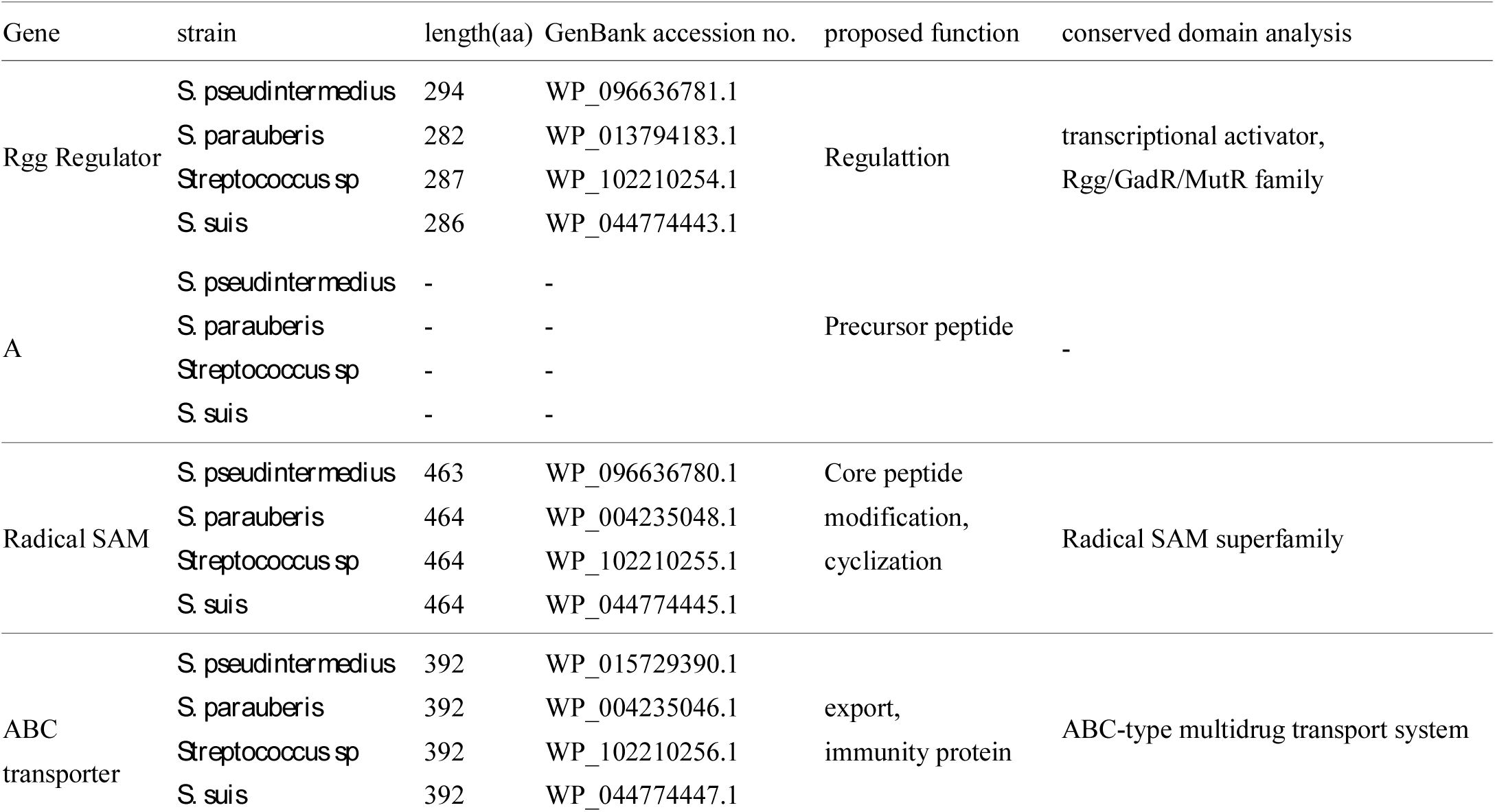
Proposed functions of the ORFs in the ITRRRY type of RiPPs biosynthetic pathways.

| Gene | strain | length(aa) | GenBank accession no. | proposed function | conserved domain analysis |
| --- | --- | --- | --- | --- | --- |
| Rgg Regulator | *S. pseudintermedius* | 294 | WP_096636781.1 | Regulation | transcriptional activator, Rgg/GadR/MutR family |
|  | *S. parauberis* | 282 | WP_013794183.1 |  |  |
|  | *Streptococcus sp* | 287 | WP_102210254.1 |  |  |
|  | *S. suis* | 286 | WP_044774443.1 |  |  |
| A | *S. pseudintermedius* | - | - | Precursor peptide | - |
|  | *S. parauberis* | - | - |  |  |
|  | *Streptococcus sp* | - | - |  |  |
|  | *S. suis* | - | - |  |  |
| Radical SAM | *S. pseudintermedius* | 463 | WP_096636780.1 | Core peptide modification, cyclization | Radical SAM superfamily |
|  | *S. parauberis* | 464 | WP_004235048.1 |  |  |
|  | *Streptococcus sp* | 464 | WP_102210255.1 |  |  |
|  | *S. suis* | 464 | WP_044774445.1 |  |  |
| ABC transporter | *S. pseudintermedius* | 392 | WP_015729390.1 | export, immunity protein | ABC-type multidrug transport system |
|  | *S. parauberis* | 392 | WP_004235046.1 |  |  |
|  | *Streptococcus sp* | 392 | WP_102210256.1 |  |  |
|  | *S. suis* | 392 | WP_044774447.1 |  |  |

## 3 Discussion

Microbial natural products are still a rich source for novel lead compounds in the treatment of many diseases[4]. Among them, RiPPs are a rapidly growing class of natural products pervasive in all domains of life[42, 43]. With the progress in genome sequencing and genome mining technology, research on the RiPPs has rapidly increased in the past few years[44]. Currently, most of the genome mining strategy for RiPPs is based on protein domain homology. Nevertheless, because the precursor peptide sequences are different and the novel tailoring enzymes could be incorporated in similar biosynthetic pathways, the newly identified RiPP gene clusters based on domain homology still prove to be fruitful in discovering new medicines[18]. Although these traditional approaches are very useful, there are a few limitations. The algorithm developed for these approaches are based on the conserved domains of the known biosynthetic enzymes. The explicitly defined rules on what is considered a new biosynthetic gene clusters in these software limits the applications and the newly discovered results are similar to known biosynthetic pathways [45]. A new structural class of compounds with a novel scaffold could not be found in most cases.

To discover a completely new family of natural products, currently the main strategy is still the screening and isolation of novel compounds[4, 7]. In these cases, after the identification of metabolites, it is still a challenge to relate the final products to their respective biosynthetic pathways. Although, diverse genome mining strategies have been developed, the regulatory patterns in the biosynthetic pathway were less studied in the past. It is reasonable to assume that different biosynthetic gene clusters might be co-regulated and common regulatory patterns exist in different clusters[45, 46]. It is important to understand the complex regulation of secondary metabolites in bacteria, especially in Streptomyces[47, 48]. It has been long known that most of the gene clusters in Streptomyces are silent under normal laboratory conditions[49, 50]. Therefore, it is necessary to study the complex regulatory networks involved in secondary metabolism and find effective ways to activate these silent gene clusters[50]. It is worth noting that regulator binding sites are often highly conserved and various tools have been developed to predict the regulator system associated with biosynthetic pathways[51]. Since the regulator sites are often conserved, it is also possible to use the regulatory sequences as a query sequence for genome mining studies. One successful example is the development of the software CASSIS for the prediction of fungal natural products present in the biosynthetic pathways[45]. In fungus, there are many gene cluster-specific regulators. Therefore, targeting the regulatory sequence could facilitate the identification of novel gene clusters.

Till date, transcription-factor centric genome-mining approach has hardly been reported for RiPPs. In this study, we investigated the genomes of streptococcal bacteria for the presence of diverse Rgg type quorum sensing system. Using this genome mining strategy, we identified more than 500 microbes having similar systems. Surprisingly, these Rgg type regulators are frequently found in conjunction with novel potential RiPP biosynthetic pathways, indicating that these quorum-sensing systems could control diverse biosynthesis of RiPP natural products. Notably, these systems are overlooked by other prediction tools based on domain homology, such as antiSMASH, since the process enzymes possess completely new domains[22, 23]. Thus, the transcription-factor centric genome-mining approach is superior to the traditional genome mining strategy that relies on protein domain similarity in this sense.

By carefully studying the downstream genes of these newly identified Rgg regulators, we found that streptococcal bacteria develop diverse strategies to produce RiPPs and most of them have not been reported previously[12]. Among all the identified systems, we chose three representatives, which had not been previously reported. Moreover, all the discovered systems are unique among all known RiPP biosynthetic gene clusters. Despite a similar “biosynthetic grammar” in all these biosynthetic pathways with a Rgg regulator, a precursor peptide, a radical SAM enzyme (or radical SAM enzyme plus a PqqD domain), and an ABC transporter, the precursor peptides and the process enzymes are completely different. A phylogenetic tree was built for all the radical SAM enzymes involved in these biosynthetic pathways. As showed in Fig. 5, the radical SAM enzymes are clearly divided into four clades. Low similarities between these process enzymes make it hard to discover these mysterious gene clusters using the traditional domain similarity-based genome mining strategy.

**Fig. 5.**
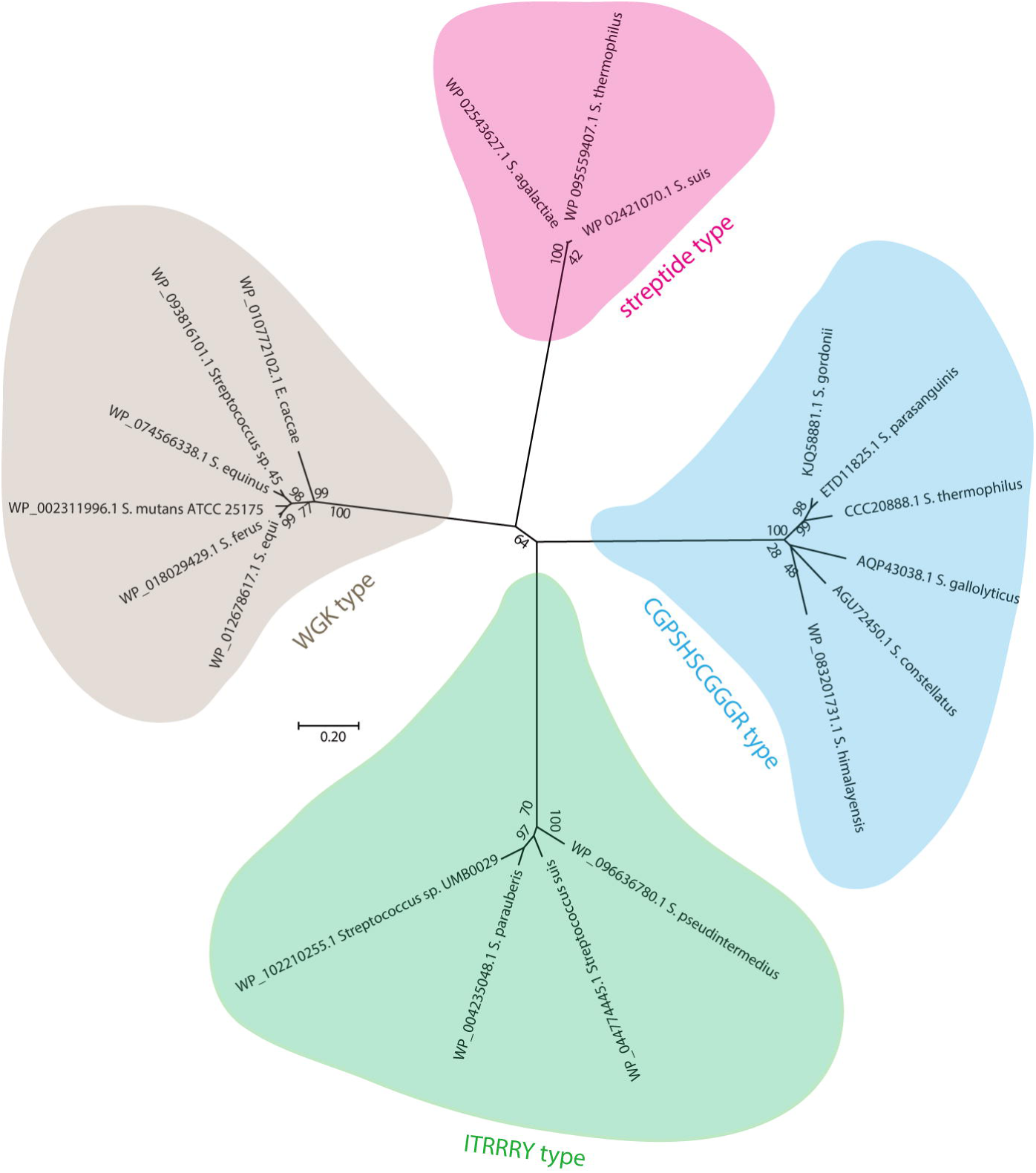
Phylogenetic tree of the Radical SAM proteins from streptide type gene clusters and the new identified gene clusters in this study.

Based on our studies, it seems that the streptococcal bacteria have developed diverse cyclization peptides system. Currently, the physiological roles of these small molecules are still unknown. It could be deduced from the streptide system that these cyclic peptides are produced at high cell densities and are triggered by the SHP like quorum sensing systems. Generally, this state is associated with virulence in the pathogenic Streptococci indicating that they might play an important role in this process[35]. On the other hand, since these systems are associated with the quorum sensing systems, it is also possible that these peptides function as self-strain activators or cross-strain inhibitors of quorum sensing-dependent pathways in Streptococci (Fig. 6)[29-31].

**Fig. 6.**
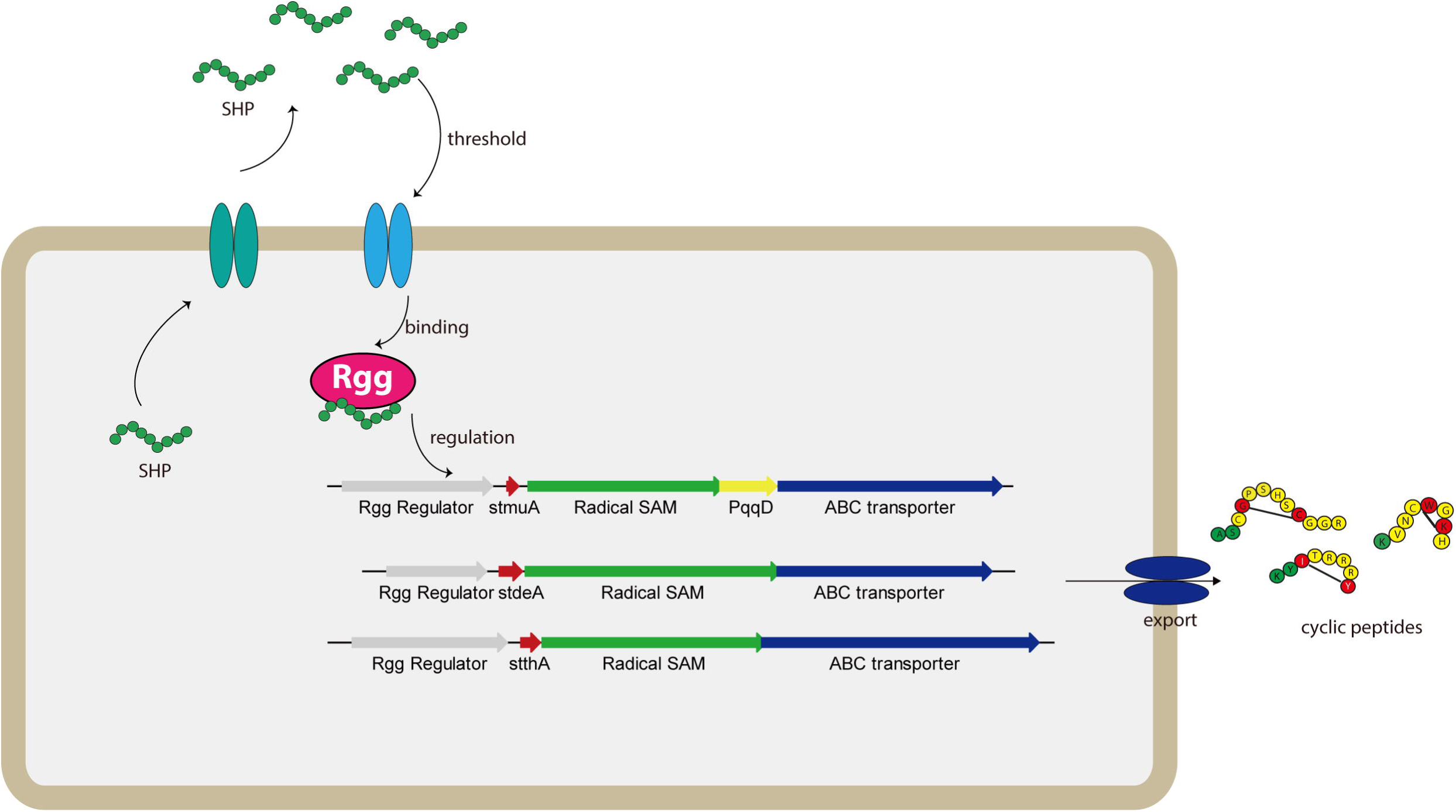
Proposed schematic of the regulation and biosynthesis of cyclic peptides in streptococcal bacteria. Rgg type of regulator control the biosynthesis of diverse crosslink peptides upon specific peptide pheromone. The peptides could be exported outside of the cell and functioned as an antibiotic or signal peptides.

## 4 Conclusion

Despite the discovery of increasing numbers of new RiPP natural products, the discovery rate of novel family of RiPPs is slow. Herein, we report several streptococcal bacteria that have the potential to produce several unique families of RiPPs, identified using a transcription-factor centric genome-mining approach. This approach could also be employed as a general and powerful tool for the discovery of other novel RiPPs. Our results also provide detailed information on these mystery biosynthetic systems. The newly identified radical SAM enzymes and their corresponding substrates provide opportunities for finding novel pathways. The detected putative RiPP biosynthetic pathways may lead to the discovery of novel compounds for the development of antibiotics. However, it is needed to point out that the identification and isolation of all these predicted cyclic peptides remains a huge challenge. Genetic and analytical methods are needed to connect the predicted genotypes to final chemotypes. Nevertheless, it is an exciting start to the identification of new molecules and studies related to these molecules are underway.

## 5 Materials and methods

### 5.1 Genome sequences

The published genome sequences of 500 bacteria (mainly streptococcal bacteria) containing the Rgg type regulator were obtained from the NCBI Refseq and draft genome repository.

### 5.2 Search for Rgg regulator containing genomes

Published genomes were checked for the presence of Rgg type regulator (similar with the streptide system) by using the web-based bioinformatic tools PSI-BLAST. The Rgg gene (WP_011681385.1) from *S. thermophilus* was selected as a query sequence. The algorithm parameters were set as follows: Maximum target sequences to display was set to 500; expect value threshold was set to 10; BLOSUM62 was chosen as the matrix for scoring parameters. Predicted gene clusters in the downstream of the regulator were assigned manually and the conserved domains were analyzed by BLAST searches. The gene clusters were classified based on the similarity of the precursor peptides.

### 5.3 Phylogenetic analysis

The sequences of the Radical SAM enzymes from each gene clusters were retrieved manually from the NCBI database (www.ncbi.nlm.nih.gov/). Alignments were performed by ClustalX with standard settings. The Alignments were then used to construct the Phylogenetic trees by MEGA7. The evolutionary history was inferred by using the Neighbor-Joining method and the Poisson correction amino acid model. The percentage of replicate trees in which the associated taxa clustered together in the bootstrap test (1000 replicates) are also shown.

## Acknowledgments

This work was supported by National Natural Science Foundation of China (NSFC, Grant No. 21706005), China Postdoctoral Science Foundation (Grant No. 2017M610747), the Fundamental Research Funds for the Central Universities (No. ZY1713) and National Great Science and Technology Projects (2018ZX09721001).

## Author contributions

SZ and GZ designed the study and wrote the paper. The authors declare that they have no conflicts of interest with the contents of this article.

## References

[1] C.A. Arias, B.E. Murray, Antibiotic-Resistant Bugs in the 21st Century — A Clinical Super-Challenge, New Engl J Med, 360 (2009) 439-443.

[2] F.R. DeLeo, M. Otto, B.N. Kreiswirth, H.F. Chambers, Community-associated meticillin-resistant Staphylococcus aureus, The Lancet, 375 (2010) 1557-1568.

[3] J.S. Weese, E. van Duijkeren, Methicillin-resistant Staphylococcus aureus and Staphylococcus pseudintermedius in veterinary medicine, Vet Microbiol, 140 (2010) 418-429.

[4] D.J. Newman, G.M. Cragg, Natural Products as Sources of New Drugs from 1981 to 2014, J Nat Prod, 79 (2016) 629-661.

[5] B. Shen, A New Golden Age of Natural Products Drug Discovery, Cell, 163 (2015) 1297-1300.

[6] A.L. Harvey, R. Edrada-Ebel, R.J. Quinn, The re-emergence of natural products for drug discovery in the genomics era, Nat Rev Drug Discov, 14 (2015) 111-129.

[7] L.L. Ling, T. Schneider, A.J. Peoples, A.L. Spoering, I. Engels, B.P. Conlon, A. Mueller, T.F. Schaberle, D.E. Hughes, S. Epstein, M. Jones, L. Lazarides, V.A. Steadman, D.R. Cohen, C.R. Felix, K.A. Fetterman, W.P. Millett, A.G. Nitti, A.M. Zullo, C. Chen, K. Lewis, A new antibiotic kills pathogens without detectable resistance, Nature, 517 (2015) 455-459.

[8] D. Boettger, C. Hertweck, Molecular Diversity Sculpted by Fungal PKS–NRPS Hybrids, ChemBioChem, 14 (2013) 28-42.

[9] J. Piel, Biosynthesis of polyketides by trans-AT polyketide synthases, Nat Prod Rep, 27 (2010) 996-1047.

[10] M. Strieker, A. Tanović, M.A. Marahiel, Nonribosomal peptide synthetases: structures and dynamics, Curr Opin Struc Biol, 20 (2010) 234-240.

[11] G.J. Williams, Engineering polyketide synthases and nonribosomal peptide synthetases, Curr Opin Struc Biol, 23 (2013) 603-612.

[12] P.G. Arnison, M.J. Bibb, G. Bierbaum, A.A. Bowers, T.S. Bugni, G. Bulaj, J.A. Camarero, D.J. Campopiano, G.L. Challis, J. Clardy, P.D. Cotter, D.J. Craik, M. Dawson, E. Dittmann, S. Donadio, P.C. Dorrestein, K.-D. Entian, M.A. Fischbach, J.S. Garavelli, U. Goransson, C.W. Gruber, D.H. Haft, T.K. Hemscheidt, C. Hertweck, C. Hill, A.R. Horswill, M. Jaspars, W.L. Kelly, J.P. Klinman, O.P. Kuipers, A.J. Link, W. Liu, M.A. Marahiel, D.A. Mitchell, G.N. Moll, B.S. Moore, R. Muller, S.K. Nair, I.F. Nes, G.E. Norris, B.M. Olivera, H. Onaka, M.L. Patchett, J. Piel, M.J.T. Reaney, S. Rebuffat, R.P. Ross, H.-G. Sahl, E.W. Schmidt, M.E. Selsted, K. Severinov, B. Shen, K. Sivonen, L. Smith, T. Stein, R.D. Sussmuth, J.R. Tagg, G.-L. Tang, A.W. Truman, J.C. Vederas, C.T. Walsh, J.D. Walton, S.C. Wenzel, J.M. Willey, W.A. van der Donk, Ribosomally synthesized and post-translationally modified peptide natural products: overview and recommendations for a universal nomenclature, Nat Prod Rep, 30 (2013) 108-160.

[13] G.L. Challis, Genome Mining for Novel Natural Product Discovery, J Med Chem, 51 (2008) 2618-2628.

[14] A.C. Letzel, S.J. Pidot, C. Hertweck, Genome mining for ribosomally synthesized and post-translationally modified peptides (RiPPs) in anaerobic bacteria, BMC Genomics, 15 (2014) 983.

[15] J.E. Velásquez, W.A. van der Donk, Genome mining for ribosomally synthesized natural products, Curr Opin Chem Biol, 15 (2011) 11-21.

[16] M. Papagianni, Ribosomally synthesized peptides with antimicrobial properties: biosynthesis, structure, function, and applications, Biotechnol Adv, 21 (2003) 465-499.

[17] T.J. Oman, W.A. van der Donk, Follow the leader: the use of leader peptides to guide natural product biosynthesis, Nature Chem Biol, 6 (2009) 9.

[18] C. Corre, G.L. Challis, New natural product biosynthetic chemistry discovered by genome mining, Nat Prod Rep, 26 (2009) 977-986.

[19] R.D. Kersten, Y.-L. Yang, Y. Xu, P. Cimermancic, S.-J. Nam, W. Fenical, M.A. Fischbach, B.S. Moore, P.C. Dorrestein, A mass spectrometry–guided genome mining approach for natural product peptidogenomics, Nature Chem Biol, 7 (2011) 794.

[20] M.O. Maksimov, I. Pelczer, A.J. Link, Precursor-centric genome-mining approach for lasso peptide discovery, P Natl Acad Sci USA, 109 (2012) 15223-15228.

[21] M. Zerikly, G.L. Challis, Strategies for the Discovery of New Natural Products by Genome Mining, ChemBioChem, 10 (2009) 625-633.

[22] K. Blin, T. Wolf, M.G. Chevrette, X. Lu, C.J. Schwalen, S.A. Kautsar, H.G. Suarez Duran, E.L.C. de Los Santos, H.U. Kim, M. Nave, J.S. Dickschat, D.A. Mitchell, E. Shelest, R. Breitling, E. Takano, S.Y. Lee, T. Weber, M.H. Medema, antiSMASH 4.0-improvements in chemistry prediction and gene cluster boundary identification, Nucleic Acids Res, 45 (2017) W36-W41.

[23] T. Weber, K. Blin, S. Duddela, D. Krug, H.U. Kim, R. Bruccoleri, S.Y. Lee, M.A. Fischbach, R. Muller, W. Wohlleben, R. Breitling, E. Takano, M.H. Medema, antiSMASH 3.0-a comprehensive resource for the genome mining of biosynthetic gene clusters, Nucleic Acids Res, 43 (2015) W237-243.

[24] C. Zong, M.J. Wu, J.Z. Qin, A.J. Link, Lasso Peptide Benenodin-1 Is a Thermally Actuated [1]Rotaxane Switch, J Am Chem Soc, 139 (2017) 10403-10409.

[25] S. Zhu, J.D. Hegemann, C.D. Fage, M. Zimmermann, X. Xie, U. Linne, M.A. Marahiel, Insights into the Unique Phosphorylation of the Lasso Peptide Paeninodin, J Biol Chem, 291 (2016) 13662-13678.

[26] S. Zhu, C.D. Fage, J.D. Hegemann, D. Yan, M.A. Marahiel, Dual substrate-controlled kinase activity leads to polyphosphorylated lasso peptides, FEBS Lett, (2016) n/a-n/a.

[27] M. Kleerebezem, Quorum sensing control of lantibiotic production; nisin and subtilin autoregulate their own biosynthesis, Peptides, 25 (2004) 1405-1414.

[28] M. Kleerebezem, L.E.N. Quadri, O.P. Kuipers, W.M. De Vos, Quorum sensing by peptide pheromones and two-component signal-transduction systems in Gram-positive bacteria, Mol Microbiol, 24 (1997) 895-904.

[29] B. Fleuchot, C. Gitton, A. Guillot, J. Vidic, P. Nicolas, C. Besset, L. Fontaine, P. Hols, N. Leblond-Bourget, V. Monnet, R. Gardan, Rgg proteins associated with internalized small hydrophobic peptides: a new quorum-sensing mechanism in streptococci, Mol Microbiol, 80 (2011) 1102-1119.

[30] R. Gardan, C. Besset, A. Guillot, C. Gitton, V. Monnet, The oligopeptide transport system is essential for the development of natural competence in Streptococcus thermophilus strain LMD-9, J Bacteriol, 191 (2009) 4647-4655.

[31] M. Ibrahim, A. Guillot, F. Wessner, F. Algaron, C. Besset, P. Courtin, R. Gardan, V. Monnet, Control of the transcription of a short gene encoding a cyclic peptide in Streptococcus thermophilus: a new quorum-sensing system?, J Bacteriol, 189 (2007) 8844-8854.

[32] K.M. Davis, K.R. Schramma, W.A. Hansen, J.P. Bacik, S.D. Khare, M.R. Seyedsayamdost, N. Ando, Structures of the peptide-modifying radical SAM enzyme SuiB elucidate the basis of substrate recognition, P Natl Acad Sci USA, 114 (2017) 10420-10425.

[33] K.R. Schramma, L.B. Bushin, M.R. Seyedsayamdost, Structure and biosynthesis of a macrocyclic peptide containing an unprecedented lysine-to-tryptophan crosslink, Nature Chem, 7 (2015) 431-437.

[34] K.R. Schramma, C.C. Forneris, A. Caruso, M.R. Seyedsayamdost, Mechanistic Investigations of Lysine-Tryptophan Cross-Link Formation Catalyzed by Streptococcal Radical S-Adenosylmethionine Enzymes, Biochemistry, 57 (2018) 461-468.

[35] K.R. Schramma, M.R. Seyedsayamdost, Lysine-Tryptophan-Crosslinked Peptides Produced by Radical SAM Enzymes in Pathogenic Streptococci, ACS Chem Biol, 12 (2017) 922-927.

[36] L. Fluhe, M.A. Marahiel, Radical S-adenosylmethionine enzyme catalyzed thioether bond formation in sactipeptide biosynthesis, Curr Opin Chem Biol, 17 (2013) 605-612.

[37] S.F. Altschul, T.L. Madden, A.A. Schaffer, J. Zhang, Z. Zhang, W. Miller, D.J. Lipman, Gapped BLAST and PSI-BLAST: a new generation of protein database search programs, Nucleic Acids Res, 25 (1997) 3389-3402.

[38] J.A. Latham, A.T. Iavarone, I. Barr, P.V. Juthani, J.P. Klinman, PqqD is a novel peptide chaperone that forms a ternary complex with the radical S-adenosylmethionine protein PqqE in the pyrroloquinoline quinone biosynthetic pathway, J Biol Chem, 290 (2015) 12908-12918.

[39] S.R. Wecksler, S. Stoll, A.T. Iavarone, E.M. Imsand, H. Tran, R.D. Britt, J.P. Klinman, Interaction of PqqE and PqqD in the pyrroloquinoline quinone (PQQ) biosynthetic pathway links PqqD to the radical SAM superfamily, Chem Commun (Camb), 46 (2010) 7031-7033.

[40] B.J. Burkhart, G.A. Hudson, K.L. Dunbar, D.A. Mitchell, A prevalent peptide-binding domain guides ribosomal natural product biosynthesis, Nat Chem Biol, 11 (2015) 564-570.

[41] S. Zhu, C.D. Fage, J.D. Hegemann, A. Mielcarek, D. Yan, U. Linne, M.A. Marahiel, The B1 Protein Guides the Biosynthesis of a Lasso Peptide, Sci Rep, 6 (2016) 35604.

[42] Manuel A. Ortega, Wilfred A. van der Donk, New Insights into the Biosynthetic Logic of Ribosomally Synthesized and Post-translationally Modified Peptide Natural Products, Cell Chem Biol, 23 (2016) 31-44.

[43] L. Liu, T. Hao, Z. Xie, G.P. Horsman, Y. Chen, Genome mining unveils widespread natural product biosynthetic capacity in human oral microbe Streptococcus mutans, Sci Rep, 6 (2016) 37479.

[44] K.J. Hetrick, W.A. van der Donk, Ribosomally synthesized and post-translationally modified peptide natural product discovery in the genomic era, Curr Opin Chem Biol, 38 (2017) 36-44.

[45] T. Wolf, V. Shelest, N. Nath, E. Shelest, CASSIS and SMIPS: promoter-based prediction of secondary metabolite gene clusters in eukaryotic genomes, Bioinformatics, 32 (2016) 1138-1143.

[46] N. Ziemert, M. Alanjary, T. Weber, The evolution of genome mining in microbes - a review, Nat Prod Rep, 33 (2016) 988-1005.

[47] M. Daniel-Ivad, S. Pimentel-Elardo, J.R. Nodwell, Control of Specialized Metabolism by Signaling and Transcriptional Regulation: Opportunities for New Platforms for Drug Discovery?, Annu Rev Microbiol, (2018). May 23. doi:10.1146/annurev-micro-022618-042458.

[48] P.A. Hoskisson, L.T. Fernández-Martínez, Regulation of specialised metabolites in Actinobacteria – expanding the paradigms, Env Microbiol Rep, 10 (2018) 231-238.

[49] D. Mao, B.K. Okada, Y. Wu, F. Xu, M.R. Seyedsayamdost, Recent advances in activating silent biosynthetic gene clusters in bacteria, Curr Opin Microbiol, 45 (2018) 156-163.

[50] M.M. Zhang, F.T. Wong, Y. Wang, S. Luo, Y.H. Lim, E. Heng, W.L. Yeo, R.E. Cobb, B. Enghiad, E.L. Ang, H. Zhao, CRISPR-Cas9 strategy for activation of silent Streptomyces biosynthetic gene clusters, Nature Chem Biol, 13 (2017) 607.

[51] S. Hiard, R. Maree, S. Colson, P.A. Hoskisson, F. Titgemeyer, G.P. van Wezel, B. Joris, L. Wehenkel, S. Rigali, PREDetector: a new tool to identify regulatory elements in bacterial genomes, Biochem Bioph Res Co, 357 (2007) 861-864.

